# Brain aerobic glycolysis and resilience in Alzheimer disease

**DOI:** 10.1101/2022.06.21.497006

**Authors:** Manu S. Goyal, Tyler Blazey, Nicholas V. Metcalf, Mark P. McAvoy, Jeremy Strain, Maryam Rahmani, Tony J. Durbin, Chengjie Xiong, Tammie L.-S. Benzinger, John C. Morris, Marcus E. Raichle, Andrei G. Vlassenko

**Affiliations:** Mallinckrodt Institute of Radiology, Washington University School of Medicine, St. Louis, MO, USA; Department of Neurology, Washington University School of Medicine, St. Louis, MO, USA; Neuroimaging Labs Research Center, Washington University School of Medicine, St. Louis, MO, USA; Knight Alzheimer Disease Research Center, Washington University School of Medicine, St. Louis, MO, USA; Department of Neuroscience, Washington University School of Medicine, St. Louis, MO, USA; Department of Biomedical Engineering, Washington University School of Medicine, St. Louis, MO, USA; Department of Psychology & Brain Science, Washington University School of Medicine, St. Louis, MO, USA

**Author notes:** Corresponding Authors, Marcus E. Raichle, Manu S. Goyal, Mallinckrodt Institute of Radiology, 510 S Kingshighway Blvd St. Louis, MO 63110.

## Abstract

The distribution of brain aerobic glycolysis (AG) in normal young adults correlates spatially with amyloid-beta (Aβ) deposition in individuals with dementia of the Alzheimer type (DAT) and asymptomatic individuals with brain amyloid deposition. Brain AG decreases with age but the functional significance of this decrease with regard to the development of DAT symptomatology is poorly understood. Using PET measurements of regional blood flow, oxygen consumption and glucose utilization—from which we derive AG—we find that cognitive impairment is strongly associated with loss of the typical youthful pattern of AG. In contrast, amyloid positivity without cognitive impairment was associated with preservation of youthful brain AG, which was even higher than that seen in typical, cognitively unimpaired, amyloid negative adults. Similar findings were not seen for blood flow nor oxygen consumption. Finally, in cognitively unimpaired adults, white matter hyperintensity burden was found to be specifically associated with decreased youthful brain AG. Our results implicate preserved AG as a factor in brain resilience to amyloid pathology and suggest that white matter disease may be a cause and/or consequence of this impaired resilience.

## INTRODUCTION

The healthy human brain largely relies upon glucose to fuel mitochondrial respiration. Yet, in young adults, a portion of resting glucose consumption exceeds that predicted by oxygen consumption rates^1^. Though the role(s) of this excess glucose utilization–i.e., aerobic glycolysis (AG), remain uncertain, some studies suggest that AG in the brain may support neurite outgrowth^2,3^, myelination^4–7^, learning^8,9^, reducing oxidative stress^10^, rapid and anticipatory neuronal activity^11^, and microglial activity^12,13^. AG in the young adult occurs more so in regions that are transcriptionally neotenous and evolutionarily expanded in humans^14^. Prior studies in humans demonstrate that brain AG decreases on average in healthy older adults, based on whole brain quantitative measurements^15,16^ as well as in terms of its regional pattern in young adults^17^. Moreover, sex influences the youthful pattern of brain glycolysis, being relatively more preserved in cognitively unimpaired aging females than in males^18^.

AG in the human brain is also affected by Alzheimer disease (AD). Whole brain estimates show that early AD is associated with a significant decrease in glucose consumption rates compared to a relatively slight change in oxygen consumption^19–21^. Interesting, amyloid deposition in both cognitively intact and impaired adults follows a regional pattern that matches that of brain AG in young adults^22,23^. Although several studies have studied total regional brain glucose consumption in relation to cognitive impairment with ^18^FDG PET, none to our knowledge has studied how regional AG is specifically affected, as compared to the larger component of brain glucose use that occurs for oxidative phosphorylation.

Here we investigate regional brain AG in AD by combining ^18^FDG PET with ^15^O-labeled H_2_O, O_2_ and CO PET to estimate glucose and oxygen metabolism together—and thereby AG—in individuals further characterized with amyloid imaging and cognitive testing. Our primary hypotheses were that youthful brain AG will be reduced in cognitively impaired individuals and preserved in cognitively intact individuals with biomarker-defined (i.e., brain amyloid positive) AD, as a reflection of brain resilience.

## RESULTS

### Study Overview

All research participants provided informed consent and all study procedures were approved by the Washington University School of Medicine Institutional Review Board. A total of 353 multi-tracer metabolic PET sessions were performed in 285 adult individuals (25-92 yo, 56% female) between the years of 2013 and 2021. Portions of these data, now labeled as the Aging Metabolism & Brain Resilience (“AMBR”) study, have been published previously^17^. Age, sex, amyloid positivity and cognitive status were defined for each individual and each of their PET imaging session(s). Amyloid status was unavailable in 80 sessions, including in only 18 individuals ≥60 yo and in 62 individuals <60 yo; note that all 40 participants <60 yo who underwent amyloid PET imaging were found to be negative. Accordingly, absent amyloid status was considered to be amyloid negative until proven otherwise. Cognitive status was typically defined using the Clinical Dementia Rating® (CDR®) scale, specifically using the sum of boxes score; when CDR could not be fully completed (n=39), cognitive status (normal versus impaired) was instead inferred from additional cognitive testing data. Further details on study procedures are provided below (see Methods).

From the metabolic PET measures, the glycolytic index (GI, a relative measure of AG), cerebral metabolic rate of glucose (CMRGlc), cerebral metabolic rate of oxygen (CMRO_2_), and cerebral blood flow (CBF) were calculated and partial volume corrected to regions defined by the Desikan-Killiany atlas and FreeSurfer subcortical parcellations. Symptomatic and “preclinical” (i.e., asymptomatic) AD were defined as individuals with brain amyloid positivity, with or without cognitive impairment, respectively.

### Analysis overview

For each PET session and independent metabolic measure (GI, CMRGlc, CMRO2, and CBF), a “youthful pattern” was defined based on its correlation to average gray matter regional values calculated in a separate, previously published but re-processed dataset comprising a cohort of young healthy adults (“N33 cohort”, 20-34 yo) (Figure 1)^24^. The N33 cohort was used to define the “youthful pattern” for each metabolic parameter in order to avoid biasing results derived from the larger AMBR cohort. However, as the N33 data were acquired approximately a decade prior with different scanner technology, an obvious outlier region (the pars orbitalis) was identified and was removed from further analysis *a priori*. The AMBR data was subjected to quantile normalization for each metabolic measure to remove “batch” effects that could have arisen during the 8 years of data collection. A Spearman rank correlation rho was then calculated for each PET session in the AMBR study as compared to the group results from the N33 cohort to calculate the “youthful index” of each metabolic measurement at the time of that PET session. These measures were subsequently related to age, sex, amyloid positivity and cognitive status using generalized linear and mixed models.

**Figure 1.**
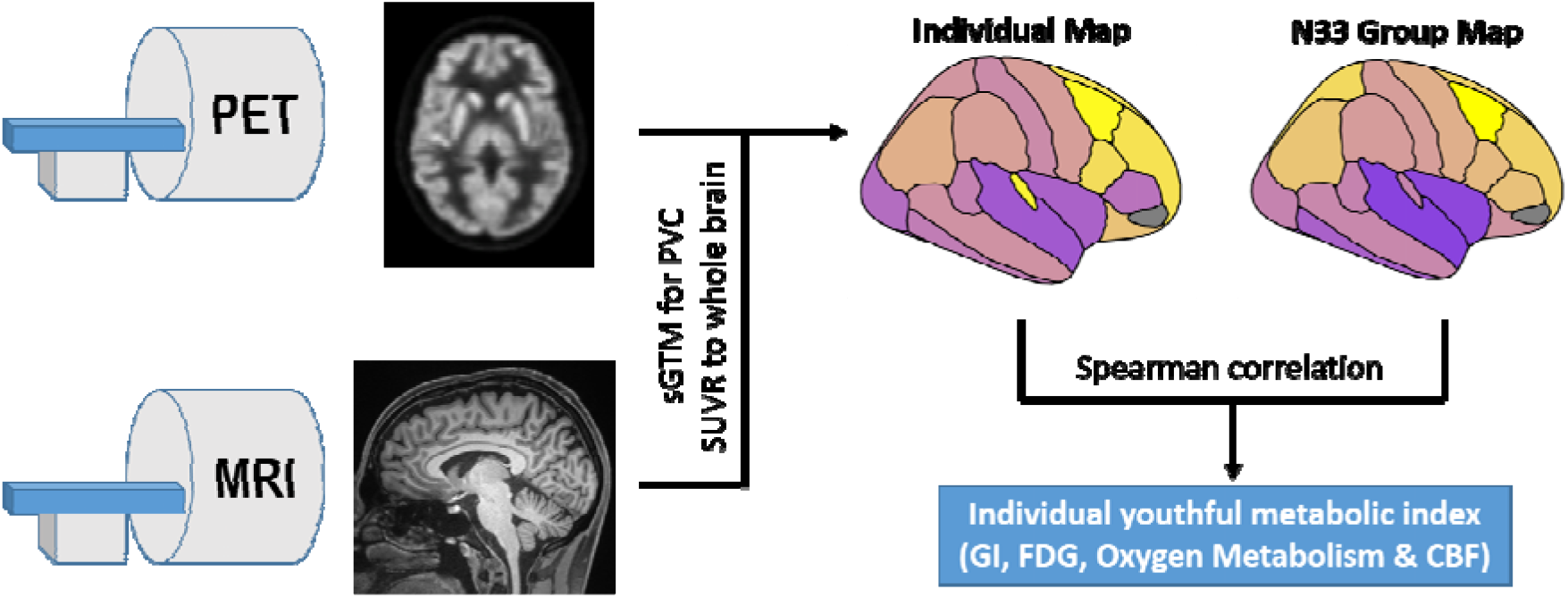
Schematic overview of youthful brain metabolism calculation. Separate resting PET and MRI scans were obtained in individuals. PET scans were preprocessed and then combined with MRI to calculate partial volume corrected regional SUVR values in the gray matter. Each individual map of brain metabolism and CBF was then compared to the corresponding group map obtained in the separate N33 young adult cohort using a Spearman correlation. The final rho value was used as the “youthful metabolic index” for that individual and specific metabolic parameter (GI, FDG, oxygen metabolism and CBF).

### Youthful brain metabolism decreases variably with age and sex

For all metabolic measures, the youthful pattern was maintained in young adults from the AMBR cohort (Figure 2). With increasing age, this youthful pattern for all metabolic parameters was variably maintained, with increasing degrees of inter-individual variability in patterns of brain metabolism, particularly for GI.

**Figure 2.**
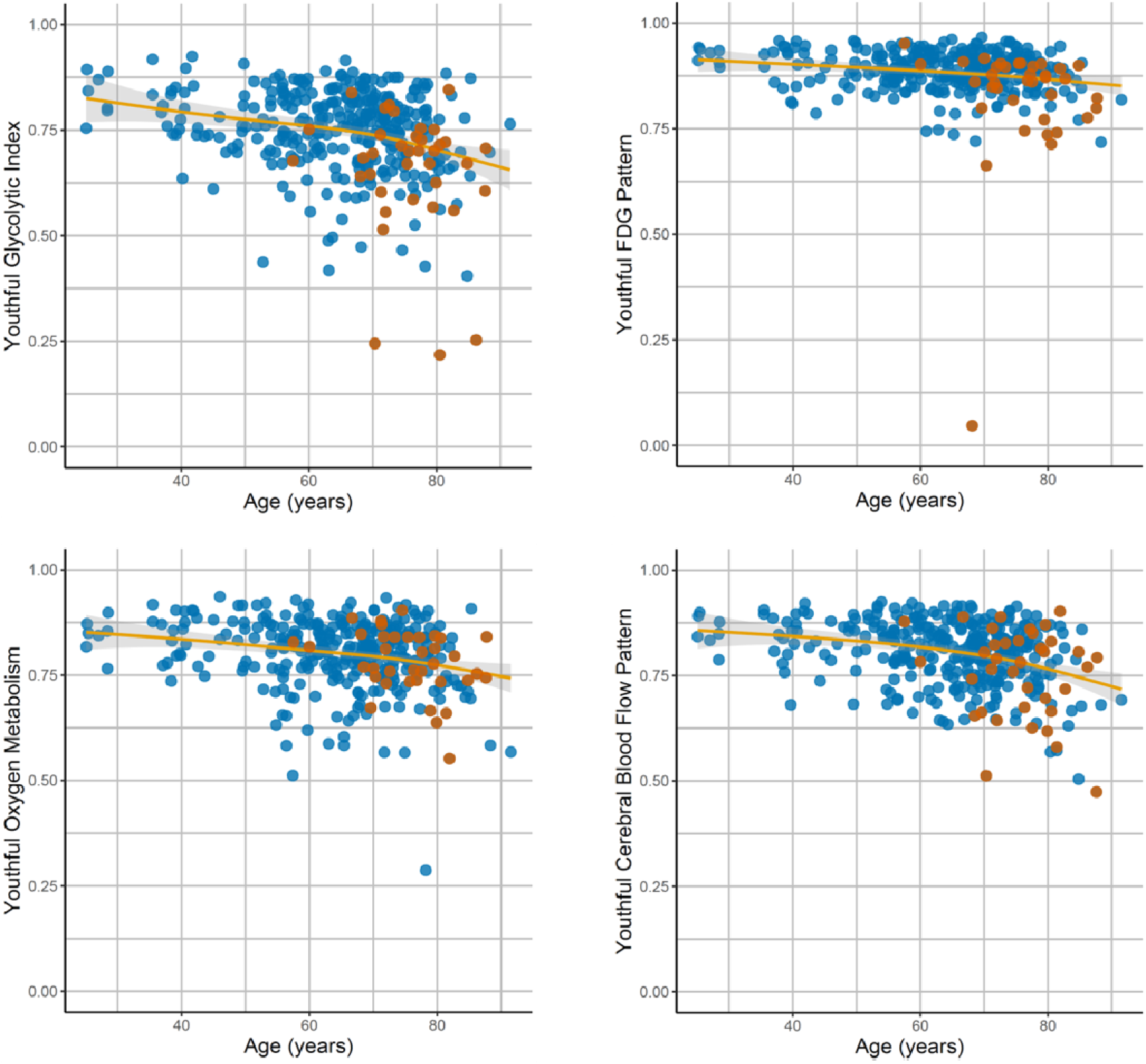
Differences in the youthful pattern of brain metabolism with age and cognitive impairment. A youthful metabolic index was calculated for aerobic glycolysis (GI), CMRGlc (FDG), oxygen metabolism (CMRO_2_) and cerebral blood flow (CBF) (see Fig. 1). This was calculated for each of the 353 PET sessions. All indices on average decreased with age (solid lines are generalized additive model [gam] fits with shaded bars reflecting standard error). However, this occurred variably, with some individuals showing a preserved youthful pattern whereas others showing a decrease in the index. Cognitively impaired individuals (red dots) were more likely to have decreased youthful GI and CMRGlc indices. This was not true, however, for CMRO2 nor CBF.

Prior observations on a subset of these data suggested that the youthful pattern of brain metabolism is more typically preserved in cognitively intact females than in males^18^. Here, female sex was again associated with a higher youthful GI index when controlling for age and amyloid status (p<0.05). This was true also for the youthful CMRO_2_ (p<0.005) and CBF (p<0.05) indices, but not significantly so for the youthful CMRGlc index. Given these findings, age and sex were included as covariates in the subsequent analyses.

### Cognitive impairment is associated with decreased youthful brain glycolysis

Cognitive impairment, as measured by CDR sum of boxes, was associated with age (p<0.005), male sex (p<0.05), and amyloid positivity (p<0.001). Cognitive impairment was further highly associated with decreased youthful GI index (p<0.0013) and youthful CMRGlc index (p<0.01), controlling for age, sex, and amyloid status. However, neither the youthful CMRO_2_ nor CBF indices were significantly associated with cognitive impairment. These results suggest early cognitive impairment is associated with changes specifically in glycolysis.

### Cognitive resilience is associated with preserved youthful brain glycolysis

If loss of the youthful glycolysis pattern during early cognitive impairment is due to direct effects of amyloid deposition rather than downstream neurodegeneration, we would predict that the youthful GI index would be lower also in amyloid positive, cognitively intact individuals. However, when restricting our analysis to individuals with CDR=0, we did not find this relationship. Instead, amyloid positivity in cognitively intact individuals correlated with *higher* youthful GI index (p<0.01), suggesting that preservation of AG in the typical glycolytic areas of youth is associated with cognitive resilience to the presence of brain amyloid (Figure 3).

**Figure 3.**
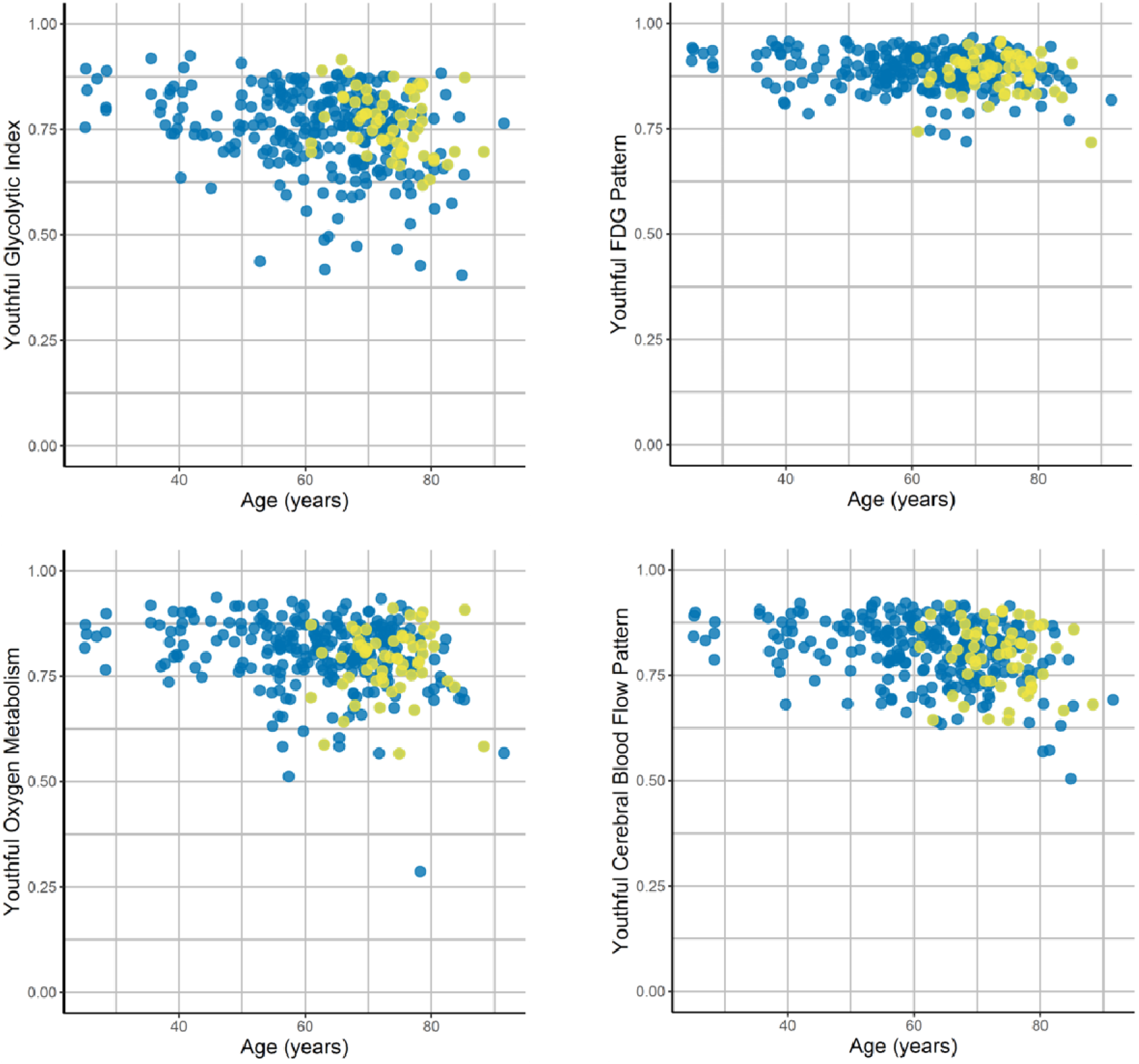
Association of amyloid positivity with youthful brain metabolism. Correlations to the youthful pattern of the GI, FDG, oxygen metabolism and cerebral blood flow were measured for each individual PET session (see Fig. 1). In this analysis only cognitively unimpaired individuals were included. Among these individuals, known amyloid positivity (yellow dots) was associated with relatively preserved youthful GI as compared to other adults, adjusting for age and sex. This was not true for CMRGlc, CMRO2 nor CBF. Blue dots reflect amyloid negativity or unknown status (n=80, including n=62 among those < 60 yo).

Given the presence of longitudinally repeated measures in a subset of individuals within the AMBR dataset, a mixed effects model was constructed to account for subjects as a random effect. The youthful GI index was further normalized using a Yeo-Johnson transformation^25^. This model again confirmed that the youthful GI index was, on average, *higher* in amyloid positive, cognitively intact individuals (p<0.05). The same analysis for total CMRGlc, CMRO_2_ and CBF did not reveal a similar significant relationship for any of these other metabolic parameters. Thus, the association between youthful brain metabolism and cognitive resilience in amyloid positive individuals is specific to brain AG.

### Specific brain regions show reduced AG in aging and AD

The results above investigate the preservation or loss of a youthful regional pattern of metabolism. This analysis was prescribed *a priori* to maximize signal-to-noise by using an omnibus measure of regional brain metabolism. However, effects of AD on brain metabolism may extend beyond a decrease in the youthful pattern of brain metabolism. We therefore computed region-by-region generalized regression models to explore the effects of age and AD on group-normalized GI within each region independently. As our metabolic data are not quantitative, only regions with significant negative changes in metabolism are included here, accounting for prior studies that demonstrate that whole brain glycolysis quantitatively decreases with age and in AD^14–16,18–20^.

Increased age was associated with decreases in GI in the superior frontal, superior parietal, caudal middle frontal, medial orbitofrontal, and entorhinal cortices and the banks of the superior temporal sulcus (Figure 4). These age related changes mirror those regions with the highest GI in young healthy adults (see Figure 1, N33 group average). Controlling for age and sex, AD status was associated with significantly reduced GI in the rostral middle frontal, inferior temporal, inferior parietal, lateral orbitofrontal, middle temporal cortices, and precuneus (Figure 4).

**Figure 4.**
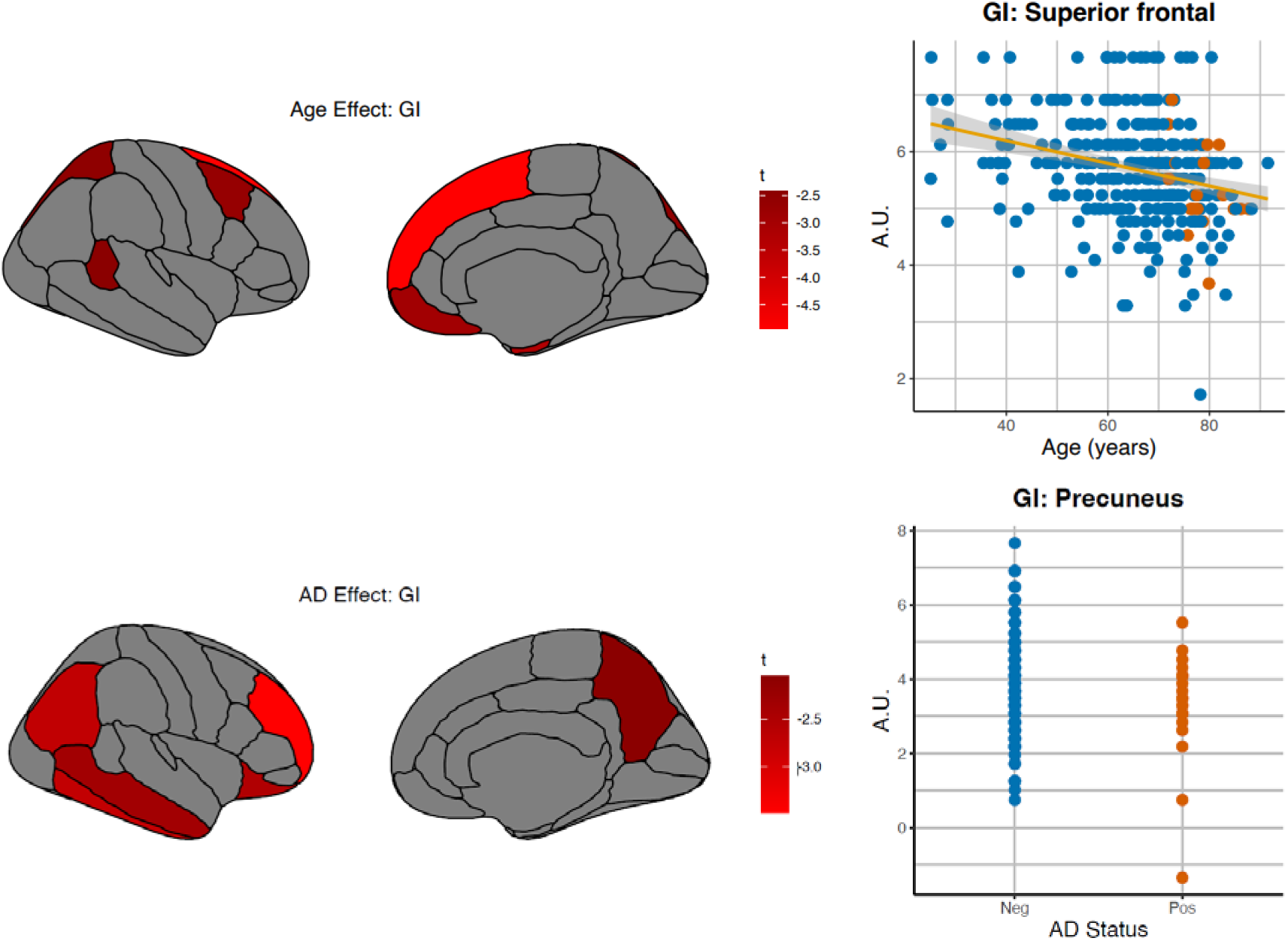
Associations of regional glycolysis (GI) with aging and Alzheimer’s disease. The associations of aging and symptomatic Alzheimer’s disease (AD) with GI were explored at a regional level. Regression models were constructed for each independent gray matter region relating GI to age, sex, and Alzheimer’s disease, and subject as a random effect. As this was an exploratory non-quantitative analysis, only regions showing a significant decrease (defined *a priori* as t-score < −1.96, uncorrected) in GI are shown here, noting that both age and AD are known to be associated with lower whole brain AG^17,19,20^. Aging was associated with relative decreases primarily in medial frontal and dorsal frontal and parietal areas, consistent with that reported previously. Alzheimer’s disease was associated with decreases in the precuneus, prefrontal, lateral parietal and temporal regions.

### White matter hyperintensity burden is specifically associated with reduced AG

White matter hyperintensities (WMH) are nearly ubiquitous in the aged human brain, though the volume of WMH varies considerably across individuals; higher volumes are associated with increased risk of cognitive decline. Given that WMH are located along tracts connecting gray matter regions, we hypothesized that increased WMH burden might be a key factor that reduces AG in the aging brain.

We acquired 1 mm^3^ isotropic FLAIR sequences in 142 cognitively unimpaired individuals of this cohort. WMH were then segmented using intensity thresholding, manual selection of lesions, re-thresholding, and quality control (see Methods below for details). Since WMH volumes fit a log-normal distribution in this cohort, they were then log transformed before comparing to brain AG and the other metabolic measures. Controlling for age, sex, and amyloid status, WMH volumes were significantly associated with a reduced youthful GI index (p<0.001) (Figure 5A). This was not true for the other metabolic measures (p>0.05; Figure 5B).

**Figure 5.**
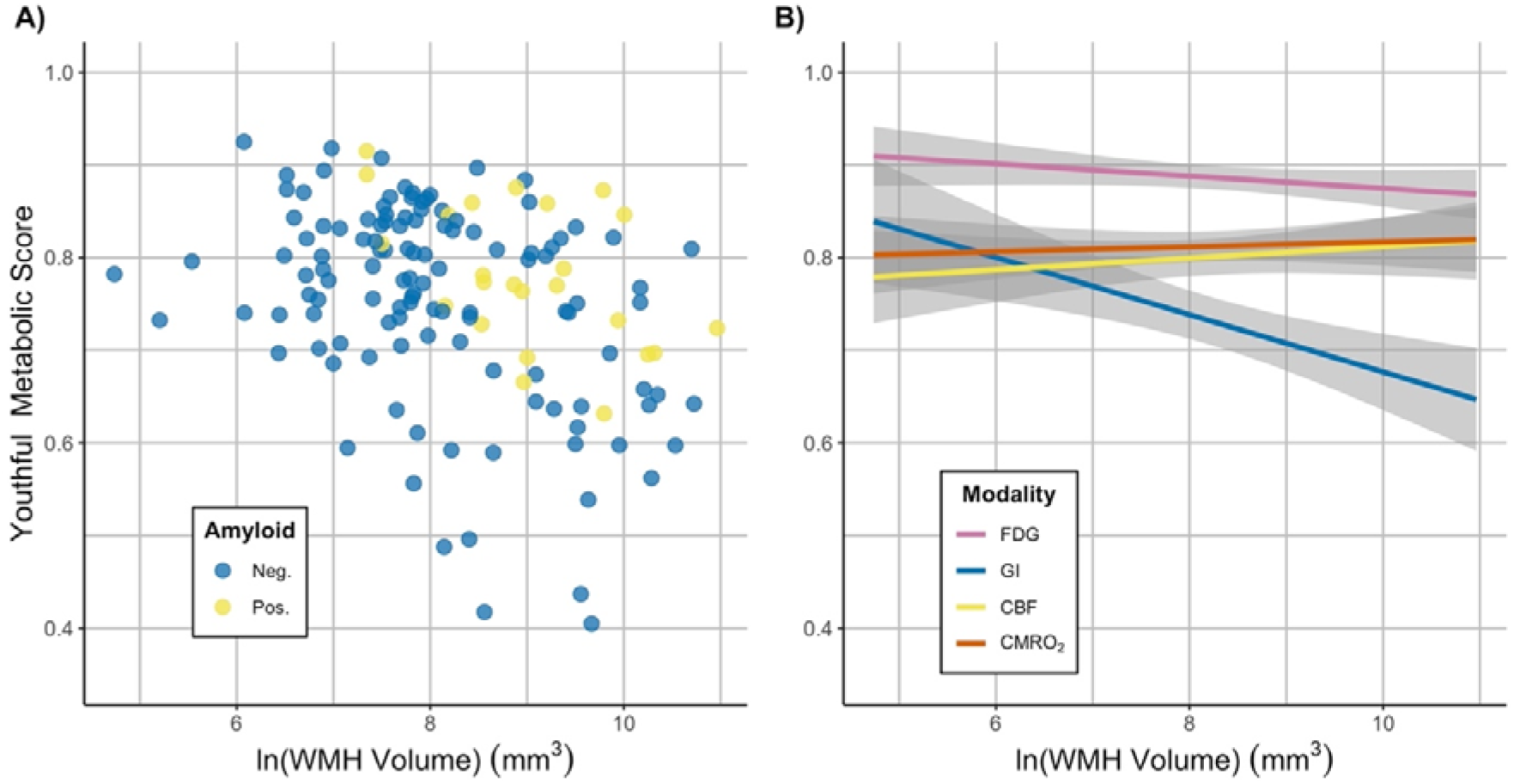
Association of white matter hyperintensities (WMH) with youthful brain metabolism in cognitively normal individuals. **A)** WMH volume (log transformed) was negatively associated with the youthful pattern of AG (GI) (*p* < 0.001). Amyloid positive participants are shown in yellow and negative individuals in blue. **B)** Model predictions for the association between youthful metabolic indices and WMH volume are shown. Unlike brain AG (“GI”, blue), neither total glucose consumption (“FDG”, pink), blood flow (“CBF”, yellow), nor oxygen consumption (“CMRO_2_”, red) was significantly associated with WMH volume (*p* > 0.05). 95% confidence intervals are shown in gray. All models in **A)** and **B)** are adjusted for age, sex, and amyloid status.

## DISCUSSION

Aging is associated with increased inter-individual variability in a variety of domains, including those related to brain function, structure, and neurodegeneration^26–28^. Here, we show that aging is similarly associated with increasingly variable changes in brain metabolism. As previously reported in a subset of these data^18^, sex accounts for some of this inter-individual variability. Now, we find that early cognitive impairment is associated with loss of the youthful pattern of brain AG and total glucose use, but not in the pattern of CMRO_2_ nor CBF. This parallels our prior PET study showing that loss of brain AG is specifically associated with tau deposition in AD^29^ as well as other whole brain studies of brain metabolism using the invasive Kety-Schmidt technique, where brain glucose consumption rates were shown to fall first during early stages of dementia, followed by decreases in oxygen consumption in later stages of dementia^19–21^. The reasons for a selective association between early symptomatic AD and loss of brain glycolysis are not yet clear, but this finding argues against ischemic processes and primary mitochondrial failure as causing a transition to early AD, since both of these would be expected to initially reduce CMRO2 and CBF and increase glycolysis. Instead, preferential loss of cells or cellular components that rely more upon glycolysis, including synapses, axons or glia, might explain why glycolysis decreases first in early AD. Loss of allostatic mechanisms and synaptic plasticity is another possible hypothesis.

In contrast to the loss of glycolysis in early dementia, asymptomatic individuals with amyloid positivity demonstrate preservation of the youthful pattern of brain AG when compared to cognitively normal individuals without amyloid pathology; a similar finding was not seen for the other metabolic parameters nor CBF. We suggest a few possible, and not mutually-exclusive, explanations for this intriguing finding. It is possible that increased AG in these individuals reflects resilience mechanisms that allow them to preserve their cognitive function despite amyloid pathology. Our results parallel a prior study showing that FDG uptake was increased in select regions of highly cognitively resilient aged individuals, including medial frontal and anterior cingulate areas, which typically show high aerobic glycolysis during youth^30^. Accordingly, much like Wald’s famous analysis of survivorship bias when investigating bullet holes in returning wartime airplanes^31^, the apparent “effects” of amyloid on brain metabolism in cognitively intact individuals might actually reflect resilience mechanisms to pathology, rather than amyloid-related damage. Cohort studies like these that require participant dedication, resources and/or altruism might further potentiate selection bias towards individuals with such resilience to neurodegeneration. These collective effects, what we have previously described as “resilience bias”^32^, might explain the relative preservation of brain AG in asymptomatic amyloid positive individuals seen here.

In vitro and animal studies have similarly shown that glycolysis is enhanced in amyloid-beta resistant neurons^33–36^. There are several potential mechanisms by which increased neuronal glycolysis might support resilience to amyloid pathology. In addition to producing NADH for oxidative capacity, glycolysis supplies several other critical metabolic pathways, including those related to the Warburg Effect, biosynthesis of lipids, nucleic and amino acids, and reducing oxidative stress, namely via the pentose phosphate pathway. Through these metabolic pathways, glycolysis might support homeostatic maintenance of neuronal networks and synaptic plasticity, which could compensate for early subclinical sites of damage and oxidative stress. Moreover, there is increasing evidence that microglial activity, which relies upon increased AG, substantially influences the brain’s response to amyloid pathology^37,38^. Animal models will be helpful to elucidate the cellular and sub-cellular locations of changes in glycolysis in the context of AD pathology.

The concept that youthful patterns of brain AG might reflect greater metabolic resilience to amyloid pathology might also help to explain why, at equivalent burdens of brain amyloid pathology, both chronological and brain age have been associated with an increased risk of cognitive impairment^39–41^. On the other hand, in young adults prior results have shown that brain amyloid preferentially deposits at sites of higher AG^22,23^; thus, the relationship between amyloid deposition and AG could well be bidirectional. Increased metabolic demand and stress might lead to failure of proteostasis, thereby leading to amyloid deposition. Conversely, amyloid deposition might incite increased metabolic stress and demand, in part as a compensatory resilience mechanism as suggested by this study^42^. Accordingly, a feed-forward loop might be established which could theoretically account for the accelerated “phase-transition” like change in amyloid deposition in the brain that is evident on longitudinal amyloid PET imaging^23^. Indeed, several studies have shown that metabolic stressors increase amyloid aggregation in the brain and even in other tissues such as pancreatic islet cells^43–45^ and transgenic *C elegans*^46^. Maintaining the supportive features of glycolysis while reducing metabolic demand might represent a means to forestall the progression of amyloid deposition worthy of further investigation.

Here we now also demonstrate that WMH burden is significantly associated with loss of the youthful pattern of gray matter brain AG. It is conceivable that WMH impact brain AG by disrupting the connectivity among gray matter regions, thereby impairing both their function and ability to maintain allostasis. Hence, WMH might be a key factor in the loss of brain resilience to pathology, including in AD where it is increasingly recognized that WMH contribute to more rapid development of dementia. Alternatively (or in addition), loss of gray matter AG might impact white matter metabolism, and thereby trigger increased vulnerability to WMH due to the well-established reliance of axons upon glycolysis from the oligodendrocyte^4,7^. Further investigations are needed to more fully understand the association(s) between WMH and brain metabolism, particularly in the white matter.

Our study has important strengths including being the largest multi-tracer metabolic PET imaging study of its kind. However, a few important limitations warrant discussion. To limit participant burden, the PET methods employed in this cohort did not include arterial lines to fully quantify measures of brain metabolism, and normalization methods were used to minimize batch effects and improve signal-to-noise, thereby limiting the inferences that can be made from these results. Future confirmatory studies using quantitative PET methods are underway, including with higher intrinsic spatial resolution and signal-to-noise to obviate the need for normalization. Another caveat in this study is that only a small number of individuals underwent longitudinal assessments; thus our results could conceivably be confounded by generational cohort effects. Ongoing longitudinal studies of brain metabolism—ideally spanning decades—would be necessary to overcome this limitation.

In conclusion, our study suggests that maintaining a youthful pattern of brain AG is associated with initial resilience to brain amyloid pathology, whereas loss of this pattern occurs alongside cognitive impairment in AD. WMH are shown to be one factor that reduces brain AG. Further research investigating mechanisms by which AG is preserved or lost in the aging brain might reveal new opportunities to improve brain resilience to pathology.

## Acknowledgments

We greatly appreciate Jennifer Byers and Kim Casey for their ongoing efforts in participant recruitment and acquiring data. We thank Abraham Z. Snyder, Matthew R. Brier and Lars Couture for their advice on the analyses of the PET and MRI imaging data. We also thank Christopher Owen in assisting in data analysis. We are particularly grateful for our research participants and their families for their altruism. We also acknowledge the directors and staff of the Neuroimaging Labs Research Center, Knight Alzheimer’s Disease Research Center, Center for Clinical Imaging Research (CCIR), and the Washington University cyclotron facility for making this research possible.

Funding for this research was provided by the Barnes-Jewish Hospital Foundation (JCM), the James S. McDonnell Foundation, the McDonnell Center for Systems Neuroscience, the NIH/NIA R01AG053503 (AGV, MER), R01AG057536 (AGV, MSG), P50AG0005681 (JCM), P01AG026276 (JCM), and P01AG003991 (JCM, TLSB). Some of the MRI sequences used to produce the AMBR dataset were obtained from the Massachusetts General Hospital. Support for Florbetapir-F18 scans at the Knight Alzheimer Disease Research Center was provided by Avid Radiopharmaceuticals, a wholly owned subsidiary of Eli Lilly.

## Author Contributions

MSG conceived of the study design, performed the analysis, and drafted and revised the manuscript. AGV conceived of the study design, oversaw data collection and processing, and revised the manuscript. TB performed and reviewed statistical analysis and revised the manuscript. NVM and MPM performed data processing and revised the manuscript. JS and MR performed data processing for WMH measurements and revised the manuscript. CX reviewed statistical analysis and revised the manuscript. MER conceived of the study design and critically revised the manuscript. JCM aided with the recruitment of participants, aided in study design, and critically revised the manuscript. TLSB aided in study design, oversaw data collection, and revised the manuscript. TJD coordinated study procedures and regulatory work, collected data, and critically revised the manuscript.

## Data and Software Availability

These data constitute the Adult Metabolism & Brain Resilience (AMBR) dataset. Data availability is based on prior subject consents and the 2018 Common Rule^47^. Coded, processed regional data prior to further data and statistical analyses are available from the study authors upon reasonable request by a qualified researcher. Further requests for raw imaging data should be directed to the VG Lab and the Knight Alzheimer Disease Research Center studies from which these data were gathered (http://adrc.wustl.edu).

## METHODS

### Participants and Regulatory Approvals

This study was performed according to the principles outlined within the Declaration of Helsinki. All participants and/or their designated healthcare power of attorney consented to participation in one or more of these studies and for ongoing data analysis, as approved and overseen by the Washington University School of Medicine Institutional Review Board and the Radioactive Drug Research Committee. Data were gathered from participants enrolled in several different studies performed by the Vlassenko/Goyal (VG) Lab in the Neuroimaging Labs Research Center and the Knight Alzheimer Disease Research Center (ADRC), both at the Washington University School of Medicine in St. Louis.

A total of 285 individuals (56% women, self-reported sex / gender) aged 25-92 years were recruited from the Washington University community and the Knight ADRC. All participants had no neurological, psychiatric, or systemic medical illness that might compromise study participation. Individuals were excluded if they had contraindications to MRI, history of mental illness undergoing treatment, possible pregnancy, or medication use that could interfere with brain function. Clinical cognitive status was assessed on the basis of the Clinical Dementia Rating (CDR)^48^, or when CDR was unavailable, cognitive unimpairment was defined based on a combination of self-report and other available global cognitive tests, preferentially the AD8 or Short Blessed Test, or as a last resort, the Montreal Cognitive Assessment (MoCA) corrected for education.

### Brain metabolism PET imaging

All participants underwent metabolic brain PET and MRI structural imaging for registration and brain structure segmentation as previously described^17^. All PET images were acquired in the eyes-closed waking state. No specific instructions were given regarding cognitive activity during scanning other than to remain awake. Briefly, ^18^F-FDG, ^15^O-O_2_, ^15^O-HO_2_, and ^15^O-CO PET scans were performed on participants in the awake, eyes closed state, and processed to yield regional maps of cerebral blood flow (CBF), cerebral oxygen consumption (CMRO_2_), total cerebral glucose metabolism (CMRGlc) and aerobic glycolysis (GI). Venous samples for plasma glucose determination were obtained just before and at the midpoint of the scan to verify that glucose levels were within normal range throughout the study, as well as to obtain blood radioactivity counts during the scan for future quantitative modeling. The PET images were blurred and resampled into the Desikan-Killiany atlas space^49^. These registrations and their corresponding transformations were performed with in-house software. Individual head movement during scanning was restricted by a thermoplastic mask. GI was defined by the residuals after spatially regressing CMRO_2_ from CMRGlc^24^.

Each individual’s GI, CMRGlc, CBF, and CMRO_2_ images were partial volume corrected to regions defined by the Desikan-Killiany atlas and FreeSurfer subcortical parcellations. SUVR values were subsequently calculated for each segmented cortical and deep gray matter regions, and scaled to have whole brain means of 1. Our routine partial volume corrected PET pipeline excludes results from the frontal and temporal poles, accumbens area and parahippocampal region as these regions are highly vulnerable to noise artifact due to their location and size. All remaining regional data were than subjected to quantile normalization across PET sessions for each metabolic parameter, to account for known and unknown “batch effects” that might have occurred since the beginning of data collection in 2013. Though this normalization procedure removes quantitative information, it effectively retains rank topography while minimizing biases arising from such “batch effects” over time, including those related to scanner or radioactive tracer variability^17,50^.

### Amyloid brain PET imaging

Research amyloid brain PET imaging was performed either with ^11^C-PIB (~12 mCi) or Florbetapir-F18 (~10 mCi), injected intravenously as a single bolus followed by 60 (^11^C-PIB) or 70 (Florbetapir-F18) minutes of brain PET imaging. PET imaging was performed on a Siemens Biograph PET/CT or HR+ scanner (Siemens/CTI, Knoxville, KY).

All available amyloid imaging underwent our in-house routine amyloid brain PET processing pipeline that included the following processing steps: framewise motion correction, registration to individual MRI T1 sequences, activity extraction within FreeSurfer v5.3 segmentations based on the Desikan-Killiany Atlas^49^, and partial volume correction using the regional spread function implemented within a geometric transfer matrix framework, as has been described in detail previously^51–53^. SUVR values were subsequently calculated for each segmented cortical and deep gray matter regions, referenced to the cerebellar gray matter (i.e., cerebellum SUVR = 1). A mean cortical SUVR (MC-SUVR) was calculated by averaging the SUVR values from prefrontal, parietal and temporal cortical regions. Unless otherwise noted, a threshold MC-SUVR ≥ 1.42 is used to define a quantitatively ‘positive’ amyloid ^11^C-PIB scan, based on previously published studies^29,54–56^.

Amyloid status was considered to remain positive after any positive PET scan. Conversely, amyloid status was considered to have been negative for the duration prior to any negative PET scan. When a research amyloid scan was unavailable, the results from a clinical amyloid scan, if available, was used instead to determine amyloid status at the time of the metabolic PET session.

### MRI imaging

MRI scans were obtained in all individuals to guide anatomic localization and identify specific gray matter regions. High-resolution structural images were acquired using 1.5T (Vision, Siemens, Erlangen, Germany) and 3T (Trio or Prisma, Siemens) scanners including a 3D sagittal T1-weighted magnetization-prepared 180° radio-frequency pulses and rapid gradient-echo (MPRAGE) sequence, with resolutions ranging from 0.8 x 0.8 x 0.8 mm to 1 x 1 x 1.3 mm. In a subset of individuals undergoing 3T MRI, 1 x 1 x 1 mm isotropic FLAIR sequences were obtained for WMH assessment.

FreeSurfer Analysis: FreeSurfer v5.3 software^49,57,58^ was used to segment the brain into well-defined cortical and subcortical, gray and white matter regions of interest (ROIs) based on individual MPRAGE MRI scans using the Desikan-Killiany and base FreeSurfer subcortical atlases. These ROIs were then used for the regional estimation of all PET metabolism parameters.

### WMH measurement

WMH severity was quantified by segmenting high signal intensity regions on individual FLAIR scans using the manually segmented intensity thresholds (MSIT) method. Each FLAIR scan was first preprocessed with tools in FSL for brain extraction^59^, bias field correction^60^ and rigid body registration^61^ to an individual’s corresponding T1 image. For segmentation, an intensity threshold of ≥1.2 standard deviation (SD) was applied with an in-house MATLAB script at each axial slice. This threshold has shown to maximize the sensitivity for manually identifying WMH, as applied in other neurodegenerative cohorts^62,63^.

Manual tracings were then performed where needed by identifying true lesions from false positives due to motion, fat signal, ventricles or other sources that would not be considered WMH. To ensure that all WMH clusters were fully represented, the manually selected clusters were treated as seed regions and allowed to expand one voxel outwards all directions with a signal intensity restriction of ≥0.5 SD. All WMH binary masks were drawn by the same two raters. A neuroradiologist subsequently reviewed the WMH segmentations for accuracy.

## Notes

### Competing Interest Statement

Andrei Vlassenko: NIH research grants R01-AG057536, R01-AG053503, RF1-AG073210
Manu Goyal: NIH research grants R01-AG057536, R01-AG053503, RF1-AG073210; Stock equity in IBM, Kyndryl, Moderna and BioNTech
Marcus Raichle: NIH research grant R01-AG053503
John Morris: NIH research grants P30-AG066444, P01-AG003991, P01-AG026276, U19-AG032438; consulting fees: Barcelona Brain Research Center (BBRC), TS Srinivasan Advisory Board, Chennai, India
Tammie Benzinger: Avid Pharmaceuticals (a subsidiary of Eli Lilly and Company) provides precursor and technology transfer to Washington University for use of Florbetapir-F18.

## References

1 Blazey, T., Snyder, A. Z., Goyal, M. S., Vlassenko, A. G. & Raichle, M. E. A systematic meta-analysis of oxygen-to-glucose and oxygen-to-carbohydrate ratios in the resting human brain. PLoS One 13, e0204242, doi:10.1371/journal.pone.0204242 (2018).

2 Segarra-Mondejar, M. et al. Synaptic activity-induced glycolysis facilitates membrane lipid provision and neurite outgrowth. The EMBO journal 37, doi:10.15252/embj.201797368 (2018).

3 Chen, K. et al. Lactate transport facilitates neurite outgrowth. Bioscience reports 38, doi:10.1042/BSR20180157 (2018).

4 Funfschilling, U. et al. Glycolytic oligodendrocytes maintain myelin and long-term axonal integrity. Nature 485, 517–521, doi:10.1038/nature11007 (2012).

5 Steiner, J. et al. Clozapine promotes glycolysis and myelin lipid synthesis in cultured oligodendrocytes. Front Cell Neurosci 8, 384, doi:10.3389/fncel.2014.00384 (2014).

6 Ghosh, S., Castillo, E., Frias, E. S. & Swanson, R. A. Bioenergetic regulation of microglia. Glia 66, 1200–1212, doi:10.1002/glia.23271 (2018).

7 Nave, K. A. Myelination and the trophic support of long axons. Nat Rev Neurosci 11, 275–283, doi:10.1038/nrn2797 (2010).

8 Harris, R. A. et al. Aerobic Glycolysis Is Required for Spatial Memory Acquisition But Not Memory Retrieval in Mice. eNeuro 6, doi:10.1523/ENEURO.0389-18.2019 (2019).

9 Shannon, B. J. et al. Brain aerobic glycolysis and motor adaptation learning. Proc Natl Acad Sci U S A 113, E3782–3791, doi:10.1073/pnas.1604977113 (2016).

10 Butterfield, D. A. & Halliwell, B. Oxidative stress, dysfunctional glucose metabolism and Alzheimer disease. Nat Rev Neurosci 20, 148–160, doi:10.1038/s41583-019-0132-6 (2019).

11 Brown, A. M. & Ransom, B. R. Astrocyte glycogen as an emergency fuel under conditions of glucose deprivation or intense neural activity. Metab Brain Dis 30, 233–239, doi:10.1007/s11011-014-9588-2 (2015).

12 Baik, S. H. et al. A Breakdown in Metabolic Reprogramming Causes Microglia Dysfunction in Alzheimer’s Disease. Cell metabolism 30, 493–507 e496, doi:10.1016/j.cmet.2019.06.005 (2019).

13 Holland, R. et al. Inflammatory microglia are glycolytic and iron retentive and typify the microglia in APP/PS1 mice. Brain Behav Immun 68, 183–196, doi:10.1016/j.bbi.2017.10.017 (2018).

14 Goyal, M. S., Hawrylycz, M., Miller, J. A., Snyder, A. Z. & Raichle, M. E. Aerobic glycolysis in the human brain is associated with development and neotenous gene expression. Cell metabolism 19, 49–57, doi:10.1016/j.cmet.2013.11.020 (2014).

15 Dastur, D. K. Cerebral blood flow and metabolism in normal human aging, pathological aging, and senile dementia. J Cereb Blood Flow Metab 5, 1–9, doi:10.1038/jcbfm.1985.1 (1985).

16 Gottstein, U. & Held, K. Effects of aging on cerebral circulation and metabolism in man. Acta Neurol Scand 60, 54–55 (1979).

17 Goyal, M. S. et al. Loss of Brain Aerobic Glycolysis in Normal Human Aging. Cell metabolism 26, 353–360 e353, doi:10.1016/j.cmet.2017.07.010 (2017).

18 Goyal, M. S. et al. Persistent metabolic youth in the aging female brain. Proc Natl Acad Sci U S A 116, 3251–3255, doi:10.1073/pnas.1815917116 (2019).

19 Hoyer, S. Abnormalities of glucose metabolism in Alzheimer’s disease. Ann N Y Acad Sci 640, 53–58, doi:10.1111/j.1749-6632.1991.tb00190.x (1991).

20 Hoyer, S. Brain glucose and energy metabolism abnormalities in sporadic Alzheimer disease. Causes and consequences: an update. Exp Gerontol 35, 1363–1372, doi:10.1016/s0531-5565(00)00156-x (2000).

21 Ogawa, M., Fukuyama, H., Ouchi, Y., Yamauchi, H. & Kimura, J. Altered energy metabolism in Alzheimer’s disease. J Neurol Sci 139, 78–82 (1996).

22 Vlassenko, A. G. et al. Spatial correlation between brain aerobic glycolysis and amyloid-beta (Abeta) deposition. Proc Natl Acad Sci U S A 107, 17763–17767, doi:10.1073/pnas.1010461107 (2010).

23 Goyal, M. S. et al. Spatiotemporal relationship between subthreshold amyloid accumulation and aerobic glycolysis in the human brain. Neurobiology of Aging, doi:https://doi.org/10.1016/j.neurobiolaging.2020.08.019 (2020).

24 Vaishnavi, S. N. et al. Regional aerobic glycolysis in the human brain. Proc Natl Acad Sci U S A 107, 17757–17762, doi:10.1073/pnas.1010459107 (2010).

25 Yeo, I. K. & Johnson, R. A. A new family of power transformations to improve normality or symmetry. Biometrika 87, 954–959, doi:DOI 10.1093/biomet/87.4.954 (2000).

26 Morse, C. K. Does variability increase with age? An archival study of cognitive measures. Psychol Aging 8, 156–164, doi:10.1037//0882-7974.8.2.156 (1993).

27 Tian, Q. et al. The brain map of gait variability in aging, cognitive impairment and dementia-A systematic review. Neurosci Biobehav Rev 74, 149–162, doi:10.1016/j.neubiorev.2017.01.020 (2017).

28 Thompson, P. M. et al. Cortical variability and asymmetry in normal aging and Alzheimer’s disease. Cereb Cortex 8, 492–509, doi:10.1093/cercor/8.6.492 (1998).

29 Vlassenko, A. G. et al. Aerobic glycolysis and tau deposition in preclinical Alzheimer’s disease. Neurobiol Aging 67, 95–98, doi:10.1016/j.neurobiolaging.2018.03.014 (2018).

30 Arenaza-Urquijo, E. M. et al. The metabolic brain signature of cognitive resilience in the 80+: beyond Alzheimer pathologies. Brain 142, 1134–1147, doi:10.1093/brain/awz037 (2019).

31 Wald, A. A Reprint of’A Method of Estimating Plane Vulnerability Based on Damage of Survivors. (CENTER FOR NAVAL ANALYSES ALEXANDRIA VA OPERATIONS EVALUATION GROUP, 1980).

32 Goyal, M. S., Vlassenko, A. G. & Raichle, M. E. Reply to Biskup et al. and Tu et al.: Sex differences in metabolic brain aging. Proc Natl Acad Sci U S A 116, 10634–10635, doi:10.1073/pnas.1904673116 (2019).

33 Soucek, T., Cumming, R., Dargusch, R., Maher, P. & Schubert, D. The regulation of glucose metabolism by HIF-1 mediates a neuroprotective response to amyloid beta peptide. Neuron 39, 43–56, doi:10.1016/s0896-6273(03)00367-2 (2003).

34 Newington, J. T. et al. Amyloid beta resistance in nerve cell lines is mediated by the Warburg effect. PLoS One 6, e19191, doi:10.1371/journal.pone.0019191 (2011).

35 Lone, A., Harris, R. A., Singh, O., Betts, D. H. & Cumming, R. C. p66Shc activation promotes increased oxidative phosphorylation and renders CNS cells more vulnerable to amyloid beta toxicity. Sci Rep 8, 17081, doi:10.1038/s41598-018-35114-y (2018).

36 Arias, C., Montiel, T., Quiroz-Baez, R. & Massieu, L. beta-Amyloid neurotoxicity is exacerbated during glycolysis inhibition and mitochondrial impairment in the rat hippocampus in vivo and in isolated nerve terminals: implications for Alzheimer’s disease. Exp Neurol 176, 163–174, doi:10.1006/exnr.2002.7912 (2002).

37 Leng, F. & Edison, P. Neuroinflammation and microglial activation in Alzheimer disease: where do we go from here? Nat Rev Neurol 17, 157–172, doi:10.1038/s41582-020-00435-y (2021).

38 Knopman, D. S. et al. Alzheimer disease. Nat Rev Dis Primers 7, 33, doi:10.1038/s41572-021-00269-y (2021).

39 Schindler, S. E. et al. Predicting Symptom Onset in Sporadic Alzheimer Disease With Amyloid PET. Neurology 97, e1823–e1834, doi:10.1212/WNL.0000000000012775 (2021).

40 Millar, P. R. et al. Predicting brain age from functional connectivity in symptomatic and preclinical Alzheimer disease. Neuroimage, 119228, doi:10.1016/j.neuroimage.2022.119228 (2022).

41 Cole, J. H., Marioni, R. E., Harris, S. E. & Deary, I. J. Brain age and other bodily ‘ages’: implications for neuropsychiatry. Mol Psychiatry 24, 266–281, doi:10.1038/s41380-018-0098-1 (2019).

42 Santangelo, R. et al. beta-amyloid monomers drive up neuronal aerobic glycolysis in response to energy stressors. Aging 13, 18033–18050, doi:10.18632/aging.203330 (2021).

43 Montane, J., Klimek-Abercrombie, A., Potter, K. J., Westwell-Roper, C. & Bruce Verchere, C. Metabolic stress, IAPP and islet amyloid. Diabetes Obes Metab 14 Suppl 3, 68–77, doi:10.1111/j.1463-1326.2012.01657.x (2012).

44 Gasparini, L. et al. Effect of energy shortage and oxidative stress on amyloid precursor protein metabolism in COS cells. Neurosci Lett 231, 113–117, doi:10.1016/s0304-3940(97)00536-3 (1997).

45 Cai, Z., Zhao, B. & Ratka, A. Oxidative stress and beta-amyloid protein in Alzheimer’s disease. Neuromolecular Med 13, 223–250, doi:10.1007/s12017-011-8155-9 (2011).

46 Teo, E. et al. Metabolic stress is a primary pathogenic event in transgenic Caenorhabditis elegans expressing pan-neuronal human amyloid beta. Elife 8, doi:10.7554/eLife.50069 (2019).

47 Menikoff, J., Kaneshiro, J. & Pritchard, I. The Common Rule, Updated. N Engl J Med 376, 613–615, doi:10.1056/NEJMp1700736 (2017).

48 Morris, J. C. The Clinical Dementia Rating (CDR): current version and scoring rules. Neurology 43, 2412–2414, doi:10.1212/wnl.43.11.2412-a (1993).

49 Desikan, R. S. et al. An automated labeling system for subdividing the human cerebral cortex on MRI scans into gyral based regions of interest. Neuroimage 31, 968–980, doi:10.1016/j.neuroimage.2006.01.021 (2006).

50 Leek, J. T. et al. Tackling the widespread and critical impact of batch effects in high-throughput data. Nat Rev Genet 11, 733–739, doi:10.1038/nrg2825 (2010).

51 Su, Y. et al. Partial volume correction in quantitative amyloid imaging. Neuroimage 107, 55–64, doi:10.1016/j.neuroimage.2014.11.058 (2015).

52 Su, Y. et al. Utilizing the Centiloid scale in cross-sectional and longitudinal PiB PET studies. Neuroimage Clin 19, 406–416, doi:10.1016/j.nicl.2018.04.022 (2018).

53 Su, Y. et al. Comparison of Pittsburgh compound B and florbetapir in cross-sectional and longitudinal studies. Alzheimers Dement (Amst) 11, 180–190, doi:10.1016/j.dadm.2018.12.008 (2019).

54 Jack, C. R., Jr. et al. Age-specific and sex-specific prevalence of cerebral beta-amyloidosis, tauopathy, and neurodegeneration in cognitively unimpaired individuals aged 50-95 years: a cross-sectional study. Lancet Neurol 16, 435–444, doi:10.1016/S1474-4422(17)30077-7 (2017).

55 Mintun, M. A. et al. [11C]PIB in a nondemented population: potential antecedent marker of Alzheimer disease. Neurology 67, 446–452, doi:10.1212/01.wnl.0000228230.26044.a4 (2006).

56 Su, Y. et al. Quantitative analysis of PiB-PET with FreeSurfer ROIs. PLoS One 8, e73377, doi:10.1371/journal.pone.0073377 (2013).

57 Fischl, B. et al. Whole brain segmentation: automated labeling of neuroanatomical structures in the human brain. Neuron 33, 341–355, doi:10.1016/s0896-6273(02)00569-x (2002).

58 Fischl, B. et al. Automatically parcellating the human cerebral cortex. Cereb Cortex 14, 11–22, doi:10.1093/cercor/bhg087 (2004).

59 Smith, S. M. Fast robust automated brain extraction. Hum Brain Mapp 17, 143–155, doi:10.1002/hbm.10062 (2002).

60 Zhang, Y., Brady, M. & Smith, S. Segmentation of brain MR images through a hidden Markov random field model and the expectation-maximization algorithm. IEEE Trans Med Imaging 20, 45–57, doi:10.1109/42.906424 (2001).

61 Jenkinson, M. & Smith, S. A global optimisation method for robust affine registration of brain images. Med Image Anal 5, 143–156, doi:10.1016/s1361-8415(01)00036-6 (2001).

62 Hubbard, N. A. et al. Calibrated imaging reveals altered grey matter metabolism related to white matter microstructure and symptom severity in multiple sclerosis. Hum Brain Mapp 38, 5375–5390, doi:10.1002/hbm.23727 (2017).

63 Strain, J. et al. Depressive symptoms and white matter dysfunction in retired NFL players with concussion history. Neurology 81, 25–32, doi:10.1212/WNL.0b013e318299ccf8 (2013).

